# A perfusion-based model to explain how paclitaxel achieves tumour-selective killing

**DOI:** 10.1101/2025.09.28.679015

**Authors:** Philip Murray, Adrian Saurin

## Abstract

Paclitaxel (taxol) is a commonly used chemotherapeutic that stabilizes microtubules to inhibit chromosome segregation during mitosis. Although it is used effectively to treat a wide variety of solid tumour types, it also impairs healthy cell proliferation leading to severe dose-limiting toxicities. Newer anti-mitotic drugs have been developed but these have so far failed to offer the same clinical benefit as paclitaxel, which begs the question of why this drug targets tumour cells so effectively? Here we develop a mathematical model of paclitaxel penetration and retention within 3D tumour environments following periodic drug-on/drug-off regimes typically used in the clinic. Our model suggests that during the drug-free periods, dense poorly-perfused tissue can retain paclitaxel for much longer than well-perfused tissue. This is due to paclitaxel’s ability to bind strongly to microtubules, which causes slower drug-release from densely packed tissue. Assuming that tumour cells are generally dense and less perfused than proliferative healthy tissues, this simple model suggests that tumour-selective killing could be achieved later in each chemotherapy cycle when the drug has otherwise cleared the healthy cell compartments. We use our model to optimize dosing regimens to allow paclitaxel to selectively kill tumour spheroids of different size, whilst sparing well-perfused healthy cells. Together, our model suggests paclitaxel could target key distinguishing features of many solid tumours: their size, 3D geometry and perfusion status. It is important to validate these predictions in cell models because, if correct, they could be harnessed to optimize paclitaxel use, to predict and enhance tumour responsiveness, and to develop newer drugs that are preferentially retained for longer within solid tumours.

## Introduction

The goal of anti-cancer chemotherapy is to selectively kill tumour cells whilst minimising damage to healthy tissue. Selectivity can be achieved by targeting genetic or phenotypic features that distinguish tumour cells from healthy cells in the body. Although the goal is to develop targeted therapies that exploit genetic differences within tumour cells, most mainstream chemotherapies in use today still indiscriminately target rapidly dividing cells, a phenotypic feature of most tumour types. The problem is that various healthy cells in the body, such as immune, gut and hair follicle cells, also proliferate incredibly rapidly and are therefore also rapidly killed during chemotherapy. This causes extreme dose-limiting side-effects that ultimately limits the efficacy of treatments^1^.

Paclitaxel (also known as taxol) was originally identified in 1971^2^ and remains one of the most effective chemotherapeutics in clinical use today^3^. It stabilises microtubules to disrupt the mitotic spindle^4^, leading to either mitotic arrest or severe chromosome missegregations, both of which can cause cell death or permanent cell cycle exit^5,6^. Paclitaxel is used effectively to treat a wide range of solid tumour types, including breast, lung, prostate, ovarian, oesophageal, and bowel cancers^7–11^. However, it also causes severe life-threatening toxicities, such as neutropenia, which limit the dose, duration, and ultimately efficacy, of treatment^12^. It also affects microtubules in non-dividing cells, such as neurons, which can lead to peripheral neuropathy, another common side-effect of treatment^13^.

To overcome these limitations, the drug industry has invested billions to develop newer anti-mitotic drugs which act to specifically disrupt microtubule motors or mitotic kinases that are also essential for cell division^14^. These have proved effective at restricting tumour growth in preclinical models, but have so far been largely disappointing when compared to paclitaxel in clinical trials^15,16^. This reinforces the importance of understanding which properties of paclitaxel underpin its clinical efficacy. This could allow us to rationally design better drugs or to explain why some patients or tumour types respond better to paclitaxel than others^17^.

One major difference between paclitaxel and new generation anti-mitotic drugs relates to pharmacodynamics: when applied in 2D cell culture, paclitaxel concentrates inside cells and is released slowly upon drug washout^5,18^. This ability to become “trapped” within cells could allow paclitaxel to persist within tumours long after blood levels of the drug have declined, perhaps explaining why paclitaxel can be retained within tumours for up to a week after blood levels have fallen^19,20^. This property may be particularly important in treating human disease where tumour proliferation rates are typically much slower than in animal models.

On its own, however, retention does not explain how paclitaxel could specifically kill tumour cells in preference to healthy cells in the body. In this study, we developed a mathematical model for drug penetration and release in 3D cell environments to explore this further. Our rationale was that the dense 3D environment of most solid tumours is a major distinguishing feature that differentiates these cells from healthy cell compartments that also proliferate rapidly. Therefore, this feature could perhaps explain why paclitaxel achieves selectivity for tumour cells.

Paclitaxel uptake is known to differ markedly in spheroids compared to cell monolayers^21^. The extent of paclitaxel penetration into solid tumours increases with treatment duration and concentration^22^. The transport of paclitaxel into a solid tumor is retarded by a high tumor cell density, while the reduction in cell density mediated via drug-induced apoptosis is thought to increase penetration^22,23^. In multi-cell culture, treatment concentration must be increased by a factor of five in order to observe a cell cycle effect at depth of 150 microns^24^. Hence tissue penetration increases with treatment concentration and duration. Considering that drug penetration is restricted into deep layers in 3D cultures, we hypothesized that drug release may be similarly delayed from these later when drug levels fall. To our knowledge, the release of paclitaxel from 3D tumour masses has not been assessed or predicted previously.

To explore this further we developed a 3D model based on Chaste, an open-source software library that is used to simulate complex biological systems^25,26^. The cell-based Chaste libraries provide a suite of implementations of models of cell processes (e.g. cell cycle, division, movement, apoptosis). Chaste has been used extensively to model cell dynamics in intestinal crypts and to generate models of diffusion-limited spheroid growth. In the Chaste implementation of diffusion-limited spheroid growth, which will be used later, cells on the outer periphery experience a higher proliferation rate than those in the interior. Moreover, cells in the spheroid interior undergo hypoxia-induced cell death. Thus, in the simulations a steady state spheroid size emerges in which proliferation in the outer rim is balanced by cell loss in the hypoxic core.

We used this 3D spheroid model to examine the kinetics of paclitaxel uptake and release after drug-on and drug-off cycles typically prescribed in the clinic. Our model predicts that during the drug-free periods, paclitaxel is retained for longer within the inner cells of poorly-perfused tumour spheroids, in comparison to a layer of cells on the periphery that is directly exposed to the drug-containing media (representative of proliferative healthy cell compartments that are directly exposed to the bloodstream). This implies a fractional kill theory where tumour-selective killing is specifically achieved during the later phases of each cycle when paclitaxel has cleared the healthy cell compartments but is still retained within the tumour mass. We use our model to explore modified treatment schedules that would optimize tumour-specific killing in different sized tumour spheroids. This data implies that paclitaxel, and perhaps other anti-cancer drugs that concentrate inside cells, could target a phenotypic feature that distinguishes most solid tumours from proliferative healthy cell compartments: their dense, poor-perfused, 3D geometry.

## 1 Methods

### 1.1 Cell based model

#### 1.1.1 Paclitaxel uptake in a single cell

Let *U*(*t*) and *V*(*t*) represent the concentration of free and bound paclitaxel at time *t*, respectively. Consider dynamics given by

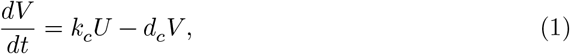

where *k*_*c*_ is the cellular uptake rate and *d*_*c*_ is an unbinding rate. Suppose that *U* (*t*) is determined by a global external forcing, i.e.

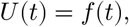

where *f*(*t*) represents the imposed concentration of paclitaxel in the blood stream determined by a clinical protocol. We consider

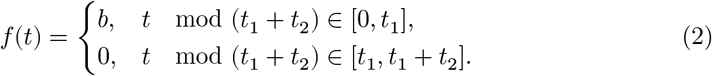

where *t*_1_ is the treatment duration, *t*_2_ is the inter-treatment duration and *b* is the treatment concentration.

#### 1.1.2 Paclitaxel uptake in fixed tissue

A fixed cell population is simulated in 2D in an approximately circular domain of radius *R*. The spatial coorindate of the *j*^*th*^ cell is given by **x**_*j*_, *j* = 1,.., *N*. Cells are positioned on a regular hexagonal lattice with unit length inter cellular separation. Free paclitaxel concentration, *u*(**x**, *t*), is represented as a continuum variable with the paclitaxel concentration of the *j*^*th*^ cell defined to be

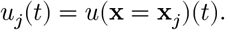

The concentration of intracellular bound paclitaxel is represented by and *v*_*j*_(*t*).

*u*(**x**, *t*) satisfies a reaction-diffusion PDE in which cells are treated as point sources/sinks,

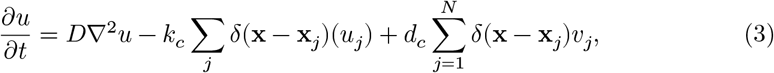

i.e. where *δ*(.) represents a Dirac delta function and *D* is the paclitaxel diffusion coefficient. A time-dependent boundary condition is imposed on the peripheral cells such that

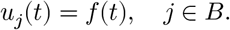

Here *B* represents the set of indices of boundary cells that reside on the periphery of the cell population.

The intracellular bound paclitaxel concentrations satisfy

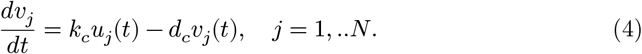

The cell-based model (Equations 3 - 4) was implemented using Chaste. Equation 3 was solved using the finite element method. To account for intracellular uptake, an intracellular regulatory network was defined in which Equation 4 was integrated for each cell.

### 1.2 A continuum model

#### 1.2.1 Model development

Representing the cell population as a continuum, a corresponding partial differential equation (PDE) model was developed that describes paclitaxel diffusion, uptake and release in a radially symmetric spheroid. Letting *u*(*r, t*) and *v*(*r, t*) represent the concentrations of free and bound paclitaxel, respectively, at radial coordinate, *r*, and time, *t*, the governing PDEs are

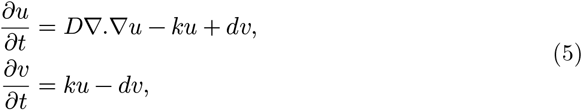

where *k* = *qk*_*c*_ and *d* = *qd*_*c*_ and *q* is the constant cell number density. A no-flux boundary condition is imposed at the origin, i.e.

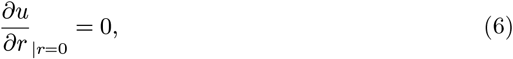

and a time-dependent Dirichlet condition on the spheroid boundary, i.e.

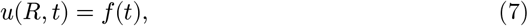

where *f*(*t*) is given by Equation 2.

#### 1.2.2 Time-averaging

Consider a solution that is time-averaged over the treatment protocol, i.e.

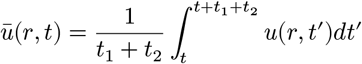

and

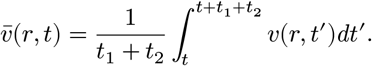

Substitution in Equations 5 - 7 yields

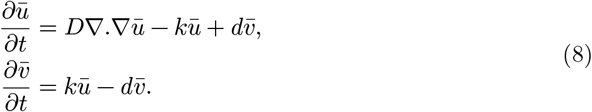

The boundary conditions are

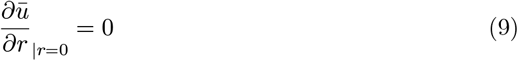

and

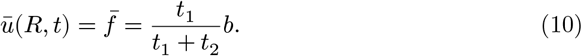

Equations 8 - 10 have a spatially homogeneous steady state given by

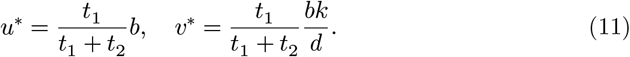

#### 1.2.3 Deviation from the time-averaged spatially homogeneous steady state

Let 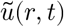 and 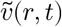 represent the deviation from the time-averaged steady state, i.e.

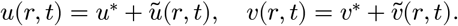

Hence Equations 5 - 7 transform to

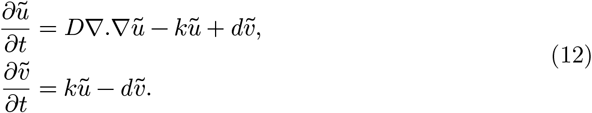

The boundary conditions are

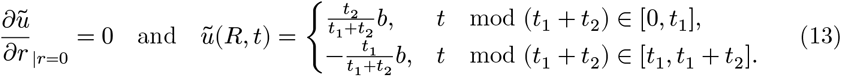

#### 1.2.4 Nondimensionalisation

We make the following scaling choices to non-dimensionalise the model:

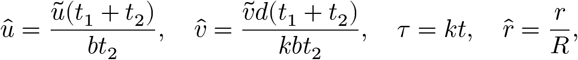

To nondimensionalise, length is scaled with the spheroid radius, *R*, time with the uptake timescale and the paclitaxel concentrations with the *on* boundary concentration. With these scalings, Equations 12 - 13 transform to

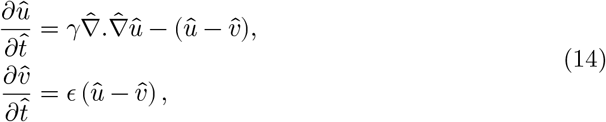

with boundary conditions

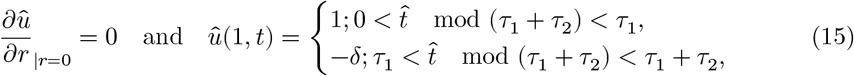

Where

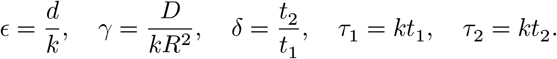

As the release rate of paclitaxel is much smaller than the uptake rate and diffusion coefficient (see Table 1)

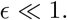

**Table 1:**
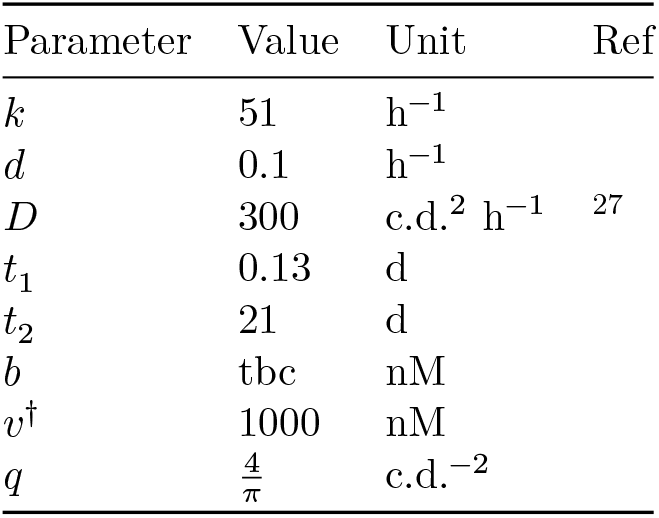
A table with model parameters used in simulations.

#### 1.2.5 A quasi-equilibrium approximation at dynamic equilibrium

Consider the limit of fast diffusion such that

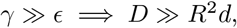

i.e. the timescale for diffusion to over the length scale of the spheroid across the spheroid is much smaller than the half-life of the bound state.

On a fast (i.e. *O*(1)) timescale

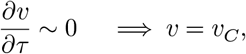

where *v*_*C*_ is a constant. Hence the governing equation for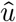 is approximated by

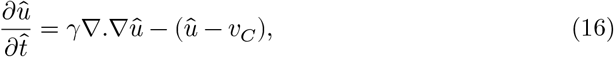

with boundary conditions

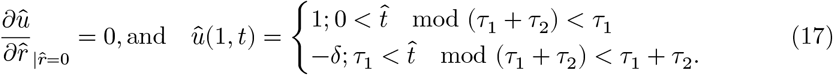

On a slow time scale (i.e. O(ε)), it is assumed that 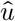 reaches quasi equilibrium and therefore satisfies the second-order modified Bessel equation

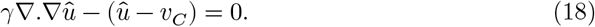

with a boundary condition that is piece-wise constant in time.

Consider the homogeneous problem

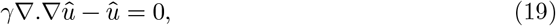

with boundary conditions

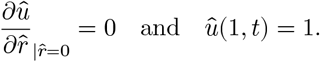

The solution is

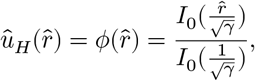

where *I*_0_(.) is a modified Bessel function of order 0.

A non-homogeneous solution to Equation 18 is

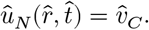

Hence a solution to Equations 17 - 18 is

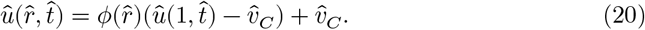

On the slow timescale 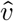 satisfies

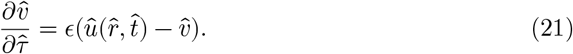

Using Equation 20 to substitute for *u* in Equation 21 yields, upon cancellation, a linear ODE at each *r*, i.e.

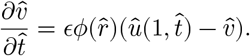

As the boundary condition 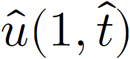 is piecewise constant, consider separately the time intervals [0, *τ*_1_] and [*τ*_1_, *τ*_2_]. Upon integration and assumption of continuity at 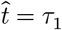,

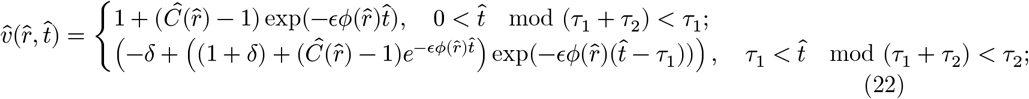

where 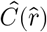 represents the local minimum of 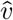, the value at the start of the cycle at (22) a given 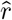.

Finally, by seeking a periodic solution at each 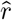 such that

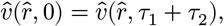

it can be shown that

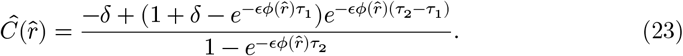

Hence Equations 22 - 23 provide a closed form periodic solution for bound paclitaxel concentration.

To obtain an upper envelope for *u*, by consider equation Equation 20 with the boundary condition 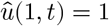 and 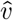 chosen to be the local minimum of the cycle *i.e*.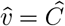.

In contrast, for the lower envelope for *u*, consider the boundary condition 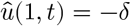 with 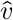 at the local maximum of the cycle *i.e*.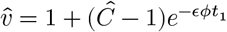.

#### 1.2.6 Protocol optimisation

To model the treatment effect on poorly perfused tissue, the bound paclitaxel concentration at the centre of a spheroid is defined to be

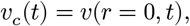

with the right-hand side computed using Equation 22 (setting *r* = 0).

For a given protocol P({*t*_1_,*t*_2_,*b*}), the fraction of a treatment cycle in which the centre of a spheroid experiences *high* bound intracellular paclitaxel is

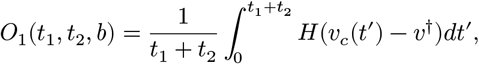

where *H*(.) is a Heaviside function defined such that

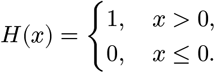

To model the treatment effect on well-perfused tissue, the bound paclitaxel concentration on the outer periphery of the spheroid is

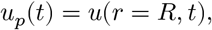

with the right-hand side computed using Equation 22 with *r* = *R*. Note that in the limit *r* → *R* we obtain the solution to Equation 1, i.e. the periphery of the spheroid exhibits dynamics described by the single cell model.

The fraction of a treatment cycle in which well-perfused cells experience paclitaxel below the critical concentration is given by

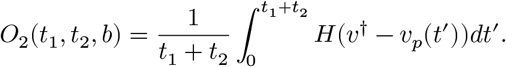

Overall, an objective function is given by

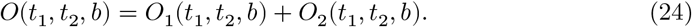

Thus, for example, a score of two represents an idealised scenario in which poorly perfused cells always experience paclitaxel concentration above the critical threshold whilst, simultaneously, well-perfused cells never do.

### 1.3 Model parameters

### 1.4 Paclitaxel uptake in a cell-based tumour spheroid model

The fixed tissue paclitaxel cellular uptake model was integrated in an existing cell based model of diffusion-limited tumour spheroid growth (^26^). Each cell has a *phase based* cell cycle model whereby cells spend prescribed amount of time in different cell cycle phases.

During M phase cell division is simulated, with a new cell being placed adjacent to the dividing cell.

Letting *c*(**x**, *t*) represent oxgen concentration. An elliptic PDE given by

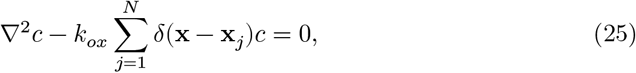

where *k*_*ox*_ is cellular uptake rate. The computed oxygen concentration is coupled to cell cycle progression such that the progression rate decreases at lower oxygen concentrations. Moreover, when oxygen falls below a hypoxic threshold, *c*_*apop*_, cells commit apoptosis. Cell mechanics are simulated by considering overdamped an equation of motion that balances cell-cell interaction forces with a damping force. Letting **x**_*i*_ represent the position of the *i*^*th*^ cell, the equations of motion takes the form

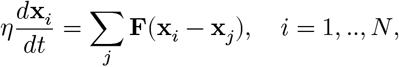

where *η* is a damping coefficent and **F** represents an intercellular force law.

A steady state spheroid size emerges in which there is balance between cell proliferation on the periphery of the spheroid and cell loss in the centre.

## 2 Results

### 2.1 Cell based model simulations

To explore paclitaxel retention in single cells, a linear model of paclitaxel cellular uptake was developed. Here paclitaxel exists in one of two states: free or intracellularly bound. Free paclitaxel binds reversibly to cellular components, with an uptake rate that is orders of magnitude larger than the unbinding rate (see Figure 1 (a)). This feature is necessary given that observed bound intracellular concentrations are orders of magnitude higher than the unbound concentrations.

**Figure 1:**
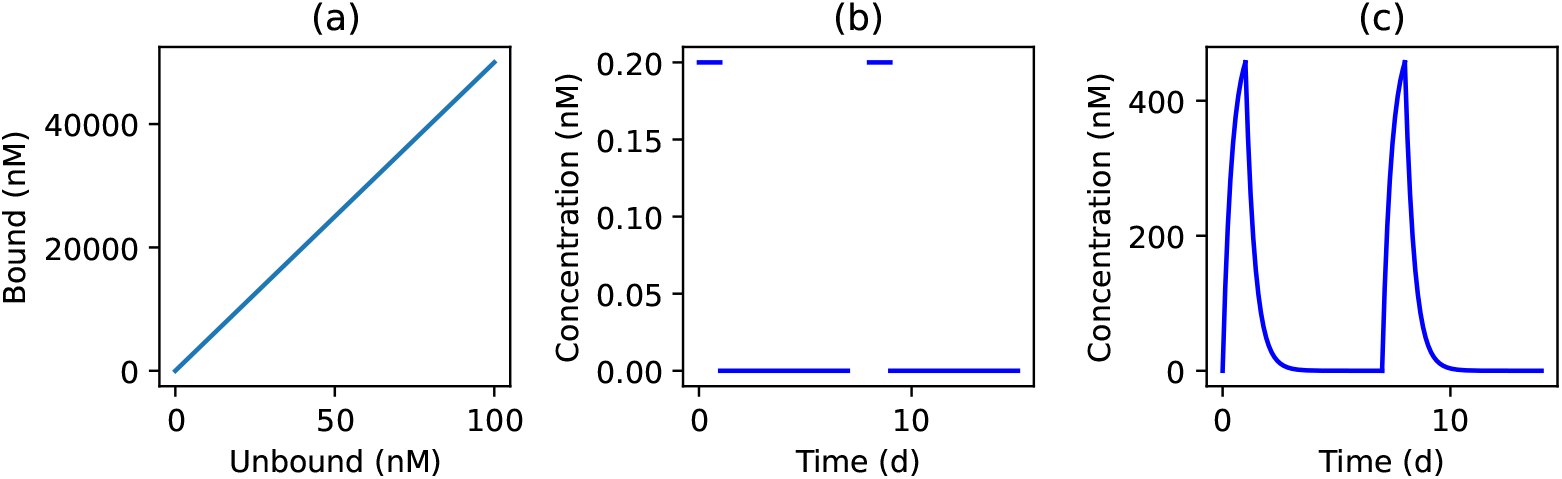
Paclitaxel concentration in a single cell model. (a) Steady state bound paclitaxel concentration is plotted against treatment concentration. (b) Periodic paclitaxel delivery protocol. (c) The bound paclitaxel concentration is plotted against time (solution of Equation 1). Here *t*_1_ = 1 d. *t*_2_ = 6.0 d, *b* = 0.2 nM. Other parameters as in Table 1.

To simulate a treatment schedule, the unbound paclitaxel concentration is modelled via a square-wave, time dependent protocol (see Figure 1 (b)). The protocol is represented by three parameters: the treatment concentration, *b*; the *on* time, *t*_1_; and the *off* time, *t*_2_. In response to this periodic stimulus, bound paclitaxel concentration increases rapidly on a time scale of minutes and decreases during the *off* part of the treatment cycle (see Figure 1 (b)).

To investigate paclitaxel uptake and retention in tissue, a cell-based model of a fixed 2D spheroid of radius *R* was developed (see Section 1.1.2). Here each cell in the simulation had a paclitaxel uptake/release model as described in Figure 1 whilst unbound paclitaxel diffused freely. Upon simulation of the cell-based model using Chaste (^25^), we observed penetration of unbound paclitaxel through the spheroid (see Figure 2a), with levels higher in the periphery than in the centre. Upon computing concentrations of bound paclitaxel, oscillation amplitude was smaller in the centre than in the periphery (see Figure 2b). Notably, the solutions appear, for the given parameter regime, to immediately find a dynamic equilibrium (i.e. fixed amplitude oscillations at a given radius).

**Figure 2:**
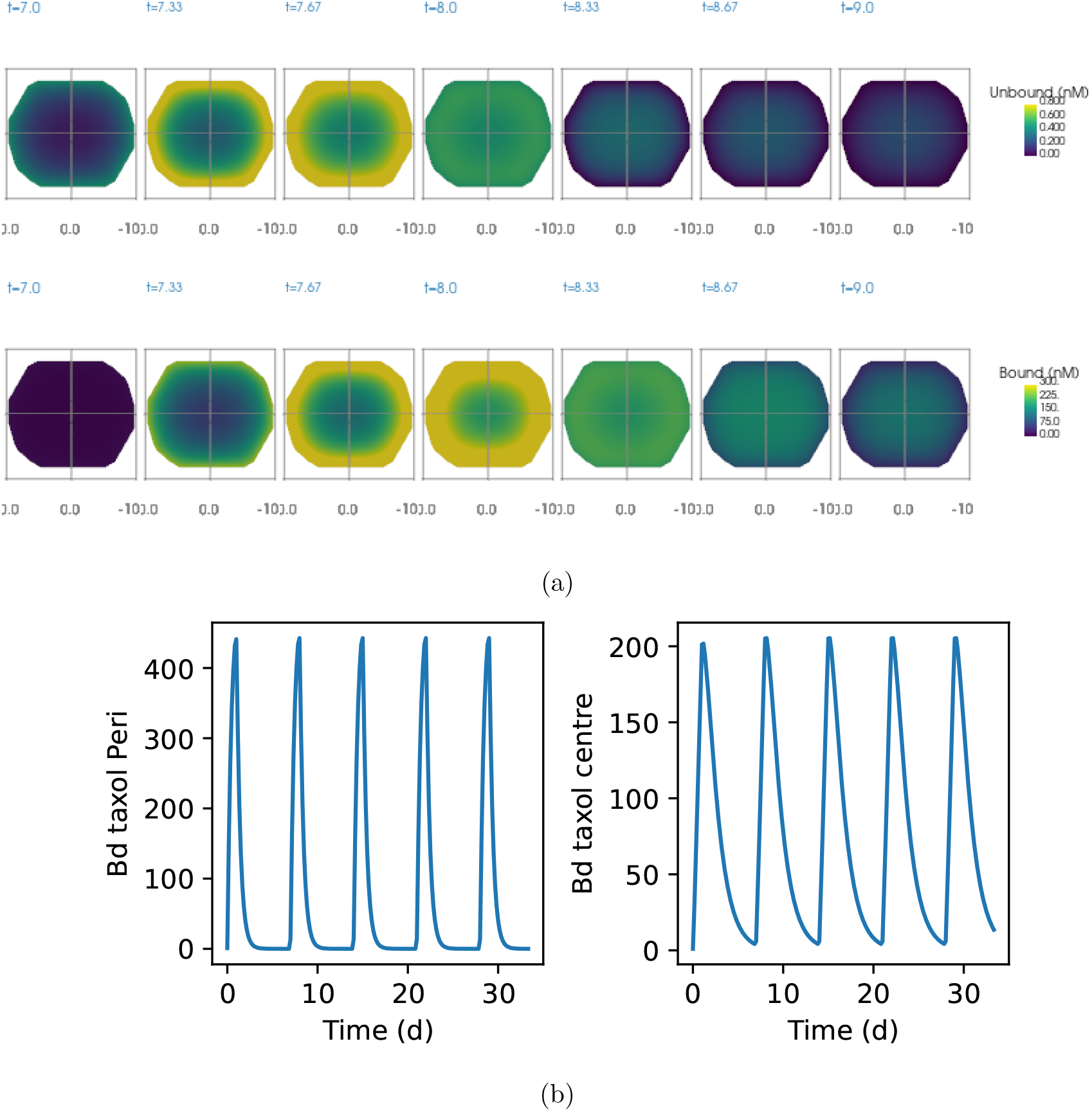
Paclitaxel uptake in a fixed tissue model under the protocol given in Figure 1 (b). (a) Snapshots of unbound (top row) and bound (bottom row) paclitaxel concentration at selected time points. (b) Bound paclitaxel concentration in the periphery and centre are plotted against time. *t*_1_ = 1. *t*_2_ = 6. Other parameters as in @#tbl-params.

Upon varying the treatment time, *t*_1_, we observed increased penetration depth (see Figure A1), behaviour that is consistent with studies of paclitaxel penetration in tumour xenografts (e.g.^22^).

To explore the effect of spheroid size on paclitaxel uptake, simulations were performed over a range of spheroid radii (see Figure 3a). Upon comparing behaviour in larger and smaller spheroids, we noted that larger spheroids had: (i) smaller amplitude oscillations of bound paclitaxel at their centre (Figure 3b); and (ii) retained paclitaxel at higher concentrations during the off part of the treatment cycle. The results suggest that well and poorly perfused regions of large spheroids experience distinct paclitaxel dynamics.

**Figure 3:**
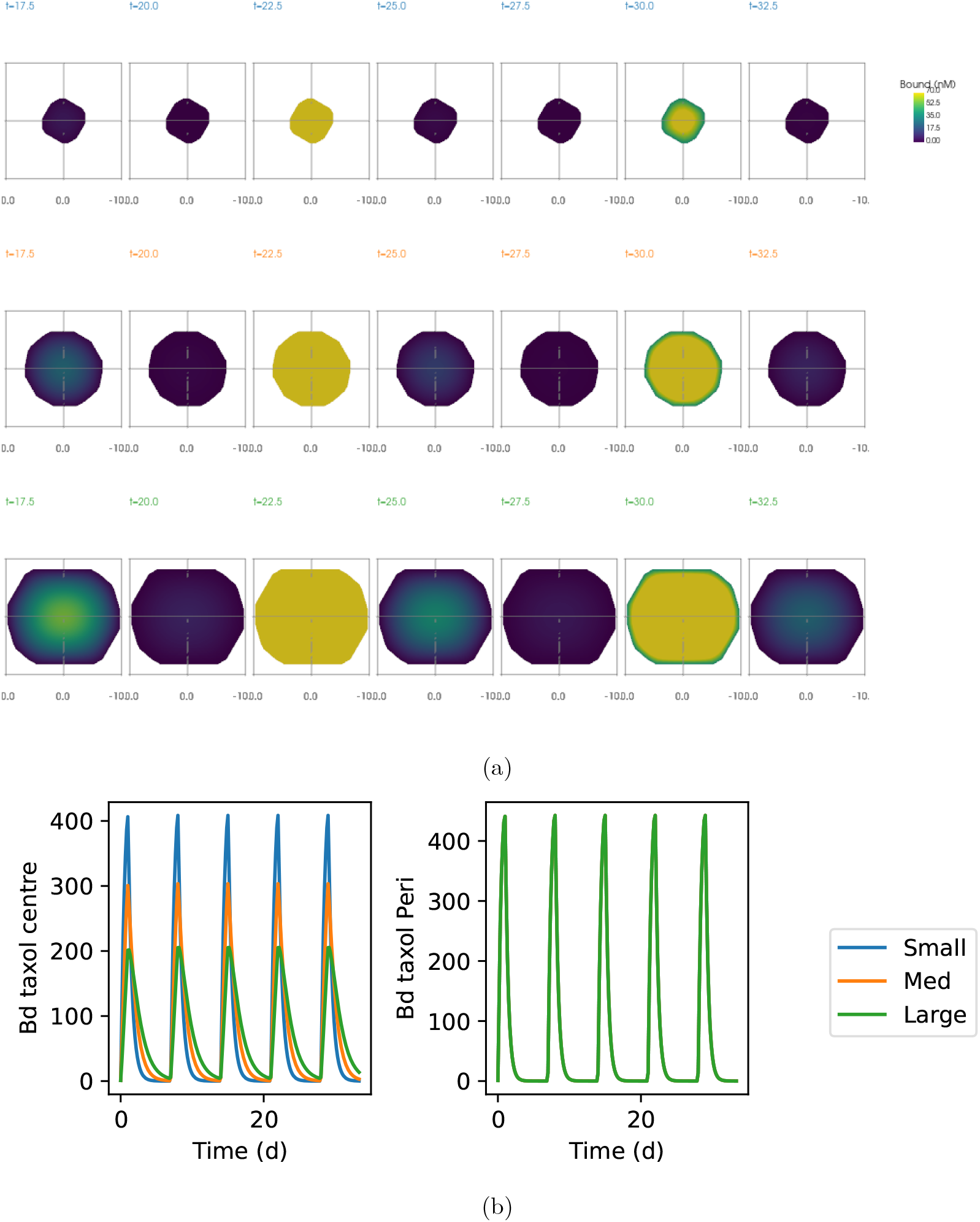
Bound paclitaxel dynamics in spheroids of different radii. (a) Top row: *R* = 4 c.d. Middle row: *R* = 8 c.d.. Bottom row: *R* = 12 c.d.. (b) Bound paclitaxel concentration is plotted against time in the centre (left) and periphery (right) of the spheroids. Other details as in Figure 2.

### 2.2 An approximation for dynamic equilibrium solution in a continuum fixed tissue model

In the fixed tissue, cell-based model presented in Figure 3b, a model for each individual cell is simulated. For simulations of large tissues run over long times, the time taken to simulate the model can quickly become prohibitive. Moreover, to infer qualitative relationships from the simulations typically requires many simulations run over different parameter values.

(b)

Hence to characterise the simulation behaviour in Figure 3b, a corresponding partial differential equation (PDE) model was developed that describes paclitaxel diffusion, uptake and release in a radially symmetric spheroid (see Section 1.2). Owing to the large disparity in uptake/release rates, a quasi-steady state approximation for unbound paclitaxel can be made (see Section 1.2.5). Hence, an approximation to the free paclitaxel concentration, *u*(*r, t*), is given by

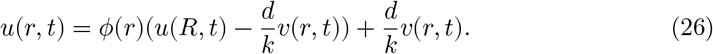

Here *ϕ*(.) is a (modified Bessel) function that decreases towards the centre of the spheroid, attaining lower values for larger spheroids (see top row in Figure 4 for plots of *ϕ* for three different sized spheroids). The envelopes for unbound paclitaxel (i.e. extreme values at a given *r*) during the on and off treatment states can also be estimated using Equation 26 (see middle row in Figure 4).

**Figure 4:**
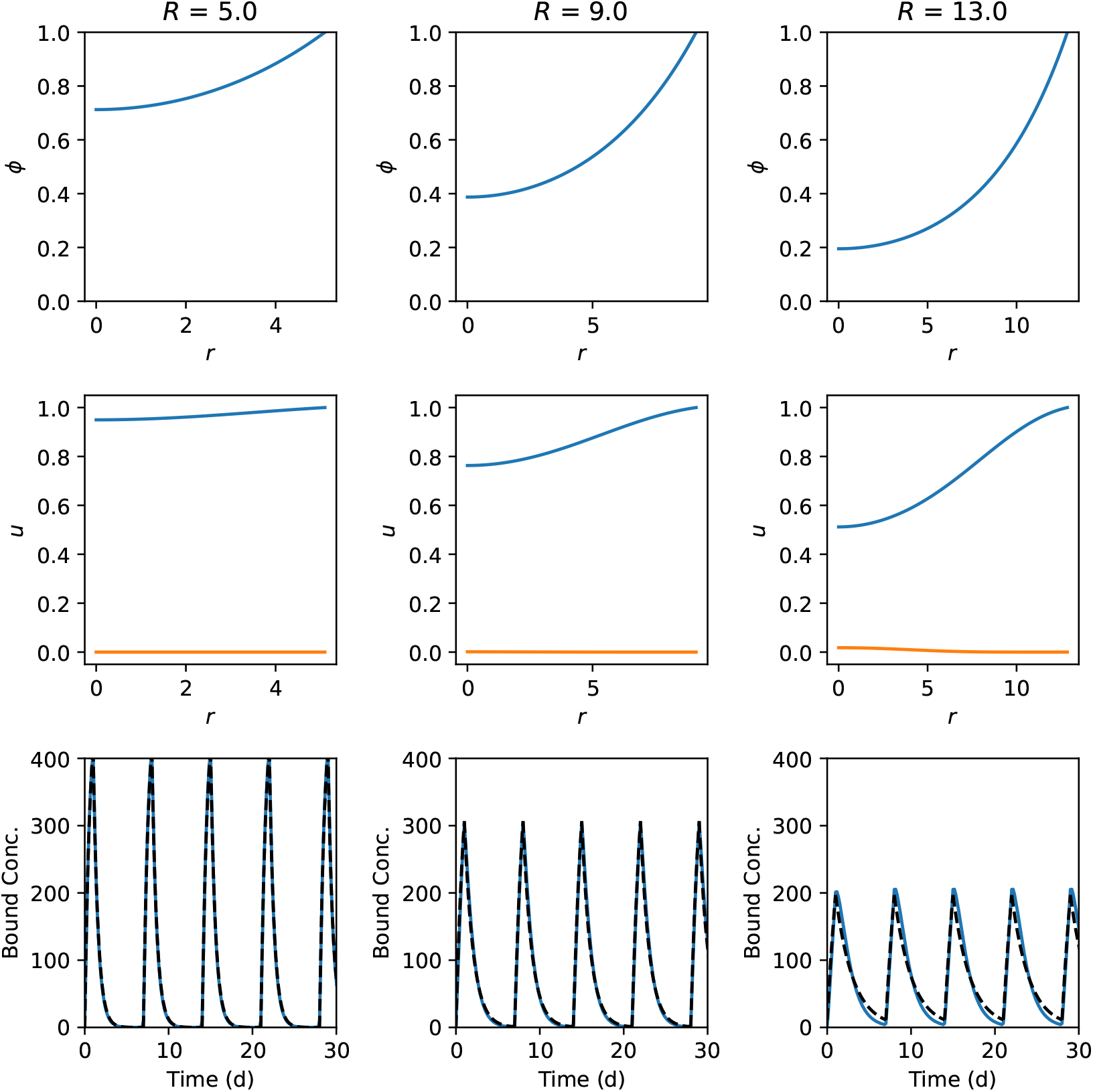
Dependence of the paclitaxel concentration on spheroid radius (columns). Top row: *ϕ*(*r*) is plotted against *r* for three different radii. Middle row: unbound paclitaxel concentration is plotted against radial coordinate, *r*, during the *on* (blue) and *off* (red) part of the treatment protocol. *u*(*r, t*) is plotted against *r*. Equation 26 is plotted for fixed *v*. Here *b* = 1.0. Bottom: bound intracellular paclitaxel concentrations in the spheroid centre are plotted against time (solid lines). Predicted dynamic equilibrium solution (Equation 28, dashed lines). Further comparisons presented in Figure A2 - Figure A4.

Using the result in Equation 26, it can be shown (see Section 1.2) that the bound paclitaxel concentration at each *r* satisfies

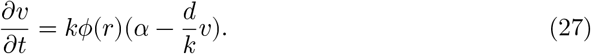

Upon integration

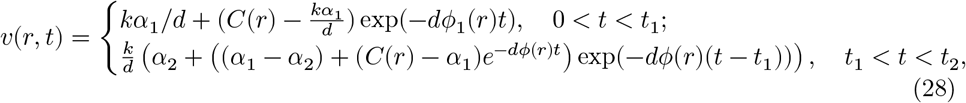

where *C*(*r*) is identified by seeking a time-periodic solution at each *r* (see Section 1.2.5).

Upon comparing the derived theoretical estimates (see Equation 28) with corresponding simulation summary statistics, we find excellent quantitative agreement (see bottom row in Figure 4). In particular, Equation 28 captures the decrease in the bound paclitaxel oscillation amplitude moving into the spheroid.

In summary, in the cell-based model simulation results paclitaxel uptake and release is differentially affected by tumour size/perfusion, with paclitaxel oscillations in the centre of large spheroids damped relative to those on the spheroid periphery. Using a quasi steady state approximation, the analytic calculations confirm this result and provide a simple interpretation: the effective paclitaxel kinetics in the centre of larger spheroids are slower those on the periphery.

### 2.3 Optimisation

We next addressed the question of whether, in the context of the developed model, an optimal treatment protocol can be identified that maximises the deleterious effect of treatment on poorly perfused tissue whilst minimising the effect on well perfused tissue.

Well-perfused tissue and poorly perfused tissue are modelled using the outer edge and centre of a large spheroid, respectively. Note that on the outer edge of the spheroid (taking the limit *r* → *R* and *ϕ* = 1) we retrieve the solution of the single cell model introduced in Section 1.1.1.

To progress, we make the modelling assumption that there is a critical bound paclitaxel concentration, *v*^†^, above which deleterious cell biological effects occur. *v* ^†^ is chosen to be of the order of micromolar concentration, as this is the lowest concentration with a clinical phenotype (it can induce mitotic spindle defects that lead to apoptosis^28^). We then defined an objective function (see Equation 24) that scored the fraction of a treatment cycle in which well perfused tissue was below the critical concentration and poorly perfused tissue was above it. Hence a score of two represents an idealised scenario where poorly perfused cells always experience paclitaxel concentration above the critical threshold whilst this never occurs for well-perfused cells.

To perform the optimisation we used a minimisation algorithm to identify values of the protocol parameters (*t*_1_,*t*_2_ and *b*) that maxmised Equation 24. This process was performed multiple times over a range of different spheroid radii, with initial values of optimisation parameters sampled uniformly, i.e.

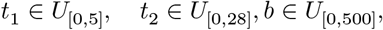

where *U* represents a uniform distribution. We found that optimal parameter sets had the following properties (Figure 5): (i) the optimal on and off times (i.e. *t* _1_ and *t*_2_) increase with spheroid radius; (ii) the optimal treatment concentration, *b*, is the maximum allowed; and (iii) the optimised value of the objective function is larger for larger spheroids (see Figure 5a), i.e. the optimal treatment schedule is more effective for larger spheroids.

**Figure 5:**
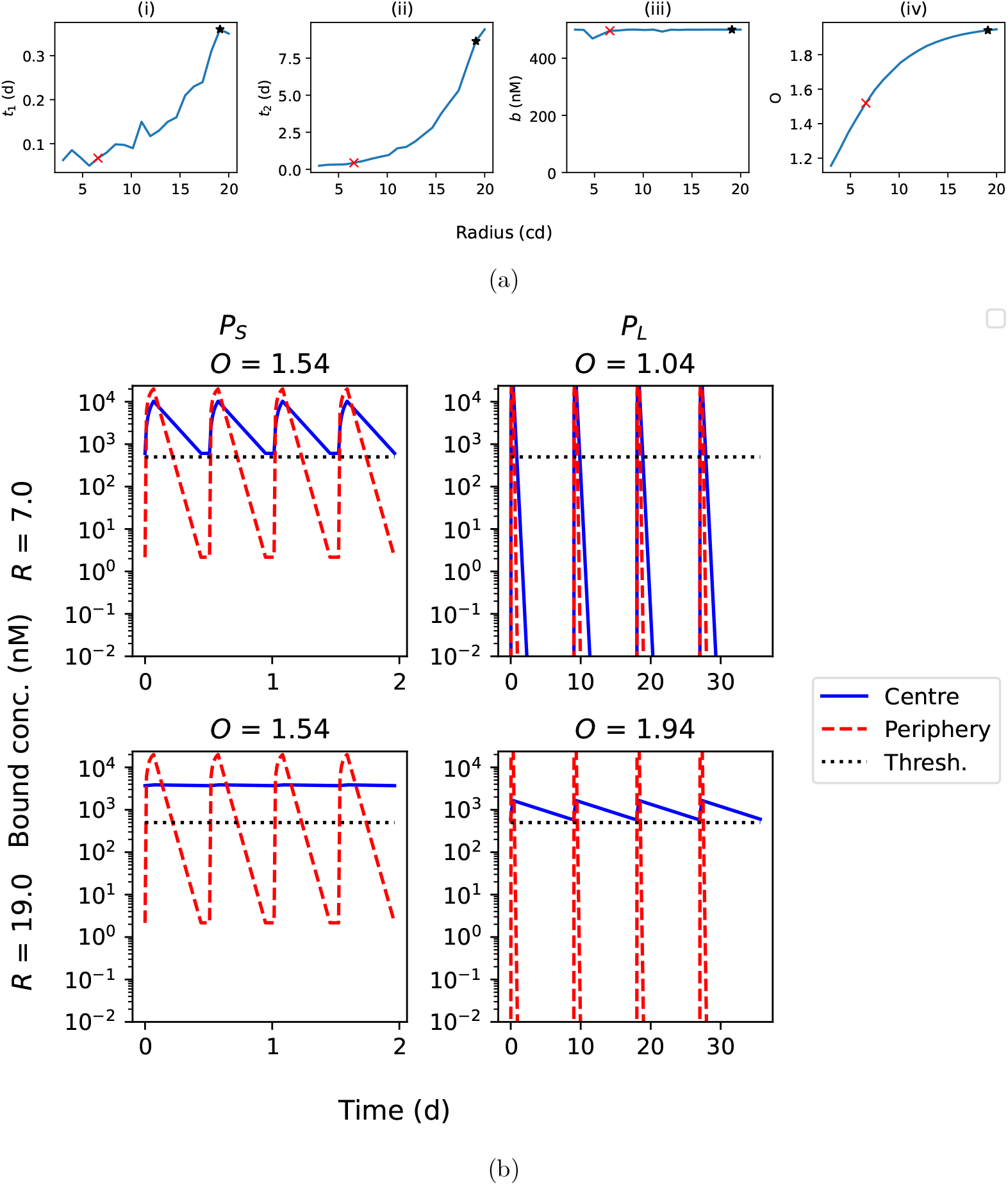
Protocol optimisation as a function of spheroid radius. (a) Top row: (i-iii) values of optimised parameters are plotted against spheroid radius. (iv) Optimised objective function is plotted against spheroid radius. Markers represent protocols optimised to small, *P*_*S*_, and large, *P*_*L*_, spheroids. (b) Bound paclitaxel dynamics for protocols optimised to small (*P*_*S*_) and large (*P*_*L*_) spheroids. Bound paclitaxel is plotted against time in spheroid centre (solid lines) and periphery (dashed lines). Critical threshold, *v*^†^ (dotted line). Optimised protocols are applied to small (top row) and large (bottom row) spheroids. See Table 2 for parameter values. Further details in Figure A5.

To illustrate model behaviour in the case of optimised protocols, two different protocols were considered that were respectively optimised to small and large spheroids (see markers in Figure 5a and Table 2). For each of the optmised protocols, we simulated the model in the case of small and large spheroids (see Figure 5b). When the protocol is matched to the spheroid size, the well-perfused cells experience minimal time with bound paclitaxel below the threshold (see diagonal entries in Figure 5b). In contrast, the centre of the spheroid experiences maximum time above the threshold.

**Table 2:** A table with parameters optimised to small/large spheroids. Parameter values taken from Figure 5b.

| Parameter/Protocol | $P_S$ | $P_L$ |
| --- | --- | --- |
| $R$ | 6.58 | 6.58 |
| $t_1$ | 0.07 | 0.36 |
| $t_2$ | 0.44 | 8.64 |
| $b$ | 496.01 | 500.0 |

Mismatching protocols results in non-optimal outcomes (see off-diagonal entries in Figure 5b). Applying a *small spheroid* protocol to a large spheroid results in the concentration of bound paclitaxel at the centre of the larger spheroids greatly exceeding the critical concentration. Here the oscillations at the centre are highly damped and the solution deviates only a little from the time-averaged steady state. In contrast, applying the *large spheroid* protocol to a small spheroid results in the concentration of bound paclitaxel falling below the critical concenration for a significant part of the treatment cycle. In both cases, applying the non-optimised protocol results in a decrease to the objective function. These results suggest that, in the context of the developed model, optimising the treatment protocol to spheroid size can have a large effect on the simulated treatment efficacy.

### 2.4 Multi-cell spheroid simulations

To investigate whether the identified paclitaxel dynamics were present in a model of spheroid growth that accounted for cell proliferation, movement and apoptosis, the paclitaxel uptake and diffusion module was integrated into the Chaste tumour spheroid model (see Section 1.4).

In the case where paclitaxel is released at apoptosis, large spheroids retain paclitaxel at higher levels than smaller spheroids (see Figure 6 for comparison of spheroids with steady state radii 4 and 8 c.d.) However, if a cell’s paclitaxel content is removed at apoptosis, we do not see a size-dependent difference in bound paclitaxel concentration at the centre of the spheroid. In this case, apoptosis acts as a sink of paclitaxel, which ultimately reduces the concentration in the centre of the spheroid.

**Figure 6:**
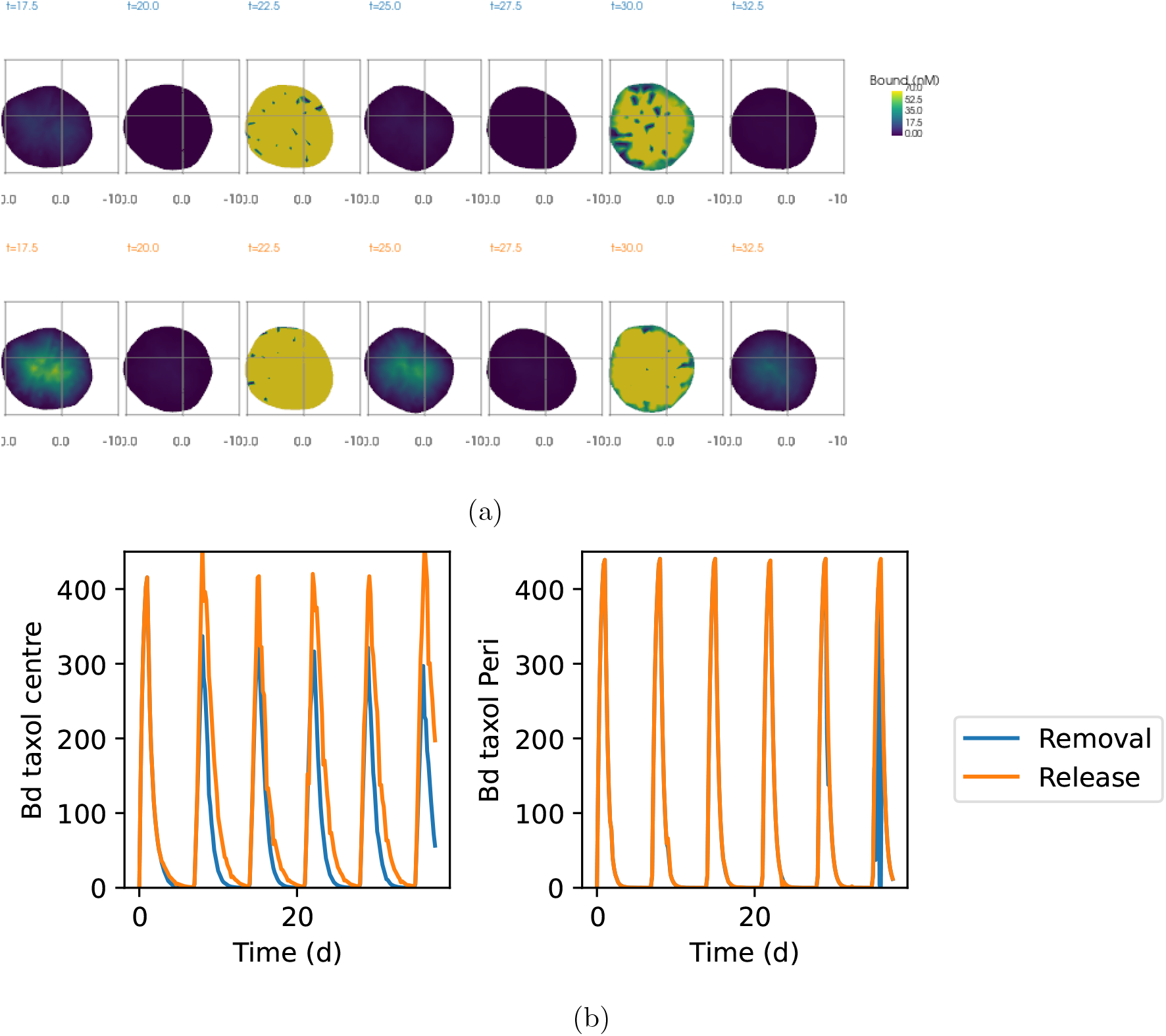
Paclitaxel uptake in a tumour spheroid model. (a) Snapshots of tumour spheroids Top: Paclitaxel is removed at apoptosis. Bottom: Paclitaxel is released at apoptosis; (b) Bound paclitaxel concentrations plotted against time in the spheroid centre (left) and periphery (right). *R* = 8 c.d. See Table 3 for additional parameter values.

## 3 Discussion

In this study we have formulated a model of paclitaxel uptake and release in tumour spheroids. A cell-based model was first considered in which paclitaxel levels were simulated in individual cells in a fixed tissue following the application of drug-on/drug-off regimens. Subsequently, we found that in the centre of larger tissues, paclitaxel concentrations had extrema that were dampened in comparison to well-perfused tissue.

**Table 3:** A table with additional model parameters used in spheroid simulation.

| Parameter | Value | Unit | Ref |
| --- | --- | --- | --- |
| $T_{G1}$ | 11 | h | |
| $T_S$ | 6 | h | |
| $T_{G2}$ | 6 | h\$ | |
| $T_M$ | 1 | h\$ | |
| $k_{spr}$ | 100 | | |
| $k_{ox}$ | 0.3 | | |
| $c_{apop}$ | 0.1 | | |

i.e. intracellular drug concentrations increased and decreased more slowly during the drug-on and drug-off periods, respectively. Hence, we reasoned that a therapeutic strategy could be developed such that treatment was optimised to maximise the effect on poorly perfused tissue whilst minimising the effect on well-perfused tissue.

To this end, we derived a corresponding PDE model in which cells were treated as a continuum. Upon making a quasi steady state approximation, an explicit formula was derived for bound intracellular paclitaxel dynamics that provided excellent agreement with the simulations. As this formula is much more efficient to compute than simulating a cell-based model, we used it to investigate treatment optimisation strategies.

An optimised treatment protocol, within the developed model framework, was defined to maximally target poorly perfused tissue whilst simultaneously minimising the effect on well-perfused tissue. The periphery of a spheroid is used as a proxy for well-perfused, healthy tissue such as intestinal epithelia. In contrast, the centre of larger spheroids is used a proxy for poorly perfused tumour cells. Treatment is represented by a periodic square wave boundary condition and characterised with three parameters: on-time, *t*_1_, off-time, *t*_2_, and treatment concentration, *b*. Moreover, it is assumed that there is some critical concentration of bound paclitaxel, *v*^†^, above which cells experience deleterious effects of treatment. Hence the goal is to identify values of *t*_1_, *t*_2_ and *b* that optimise a defined objective function.

The optimisation results indicated that maximum treatment effectiveness increases with spheroid size. Moreover, larger spheroids are better targetted with a combination of longer on and off times, with the treatment concentration being the maximum allowed value. Optimal treatments allows the centre of the spheroid to have a time-averaged value above the critical threshold. Thus cells in poorly perfused regions of tissue mainly experience deleterious effects of treatment. Meanwhile, at the spheroid periphery the time window in which cells experience high drug concentration is minimised. A web app allowing exploration of key model results can be found here.

To test if the differential paclitaxel dynamics were present in a dynamic spheroid model, the paclitaxel module was integrated into an existing cell-based model of a tumour spheroid that accounted for cell movement, proliferation and apoptosis. In this framework, the centre of larger spheroids shows accumulation of paclitaxel, so long as paclitaxel was recycled rather than lost during cell apoptosis. Thus the key feature of the fixed-tissue model upon which optimal treatment strategies rely - i.e. that the centre of larger spheroids exhibit lower amplitude oscillation - is also present upon generalising from the fixed tissue model.

In this study, protocols were optimised using fixed values of molecular parameters that have been reported in the literature (i.e. using values of the diffusion coefficient, *D*, and uptake rate, *k*). However, the degree of intratumoral accumulation of paclitaxel ranges from 9-to 172-fold across individual patients^28^. There are a range of additional tumour-specific factors that are predicted to affect paclitaxel accumulation in vivo, including tumour size and vascularization, and the makeup of the tumour stroma. In addition, our basic model assumes paclitaxel diffuses freely across the membrane, which does not act a barrier to diffusion. It also does not account for membrane transporters that can actively transport paclitaxel out of cancer cells^29^. All of these parameters would need to be reliably assessed and modelled to predict drug uptake and retention in vivo.

Nevertheless, some very basic yet important predictions can still be made from our modelling that warrant further investigation. Firstly, paclitaxel should persist within tumours in vivo for longer than it persists within healthy cell compartments. This would support a model whereby tumour-selective killing is primarily achieved later in each cycle when paclitaxel is mainly present within tumour tissue. Secondly, if some paclitaxel persists within tumours for the full duration of a treatment cycle, then there may be a concentrating effect whereby each cycle progressively increases intratumoural drug concentrations. Thirdly, the extend of paclitaxel retention within tumours may be predictive of overall treatment response. This could be assessed by measuring intratumoural concentrations during later stages of the fist chemotherapy cycle to test if these levels can be predictive of a pathological complete response (pCR). Finally, and perhaps most importantly, tumours that uptake paclitaxel at the slowest rates, will also retain the drug for the longest. This is due to the fact that microtubule binding impedes drug penetration and drug release. Therefore, strategies to selectively improve drug penetration could perhaps be used to improve overall response rates. One possibility to achieve this would be to use a modified paclitaxel that can be selectively “activated” only when it has perfused effectively into dense tumour tissue. Therefore, it can penetrate effectively without binding to micrutubules, before then being retained effectively due to microtubule binding. Various caged paclitaxel prodrugs have been developed that could be used or modified to allow this^30–32^. An alternative possibility would be to selectively promote enhanced tumour vascularization early in each cycle when the drug is highest in the bloodstream. Local hyperthermia^33^ or exercise^34^ have been shown to enhance vascularization and improve tumour drug penetration, therefore these ideas are testable within the context of controlled clinical trials. In fact, local hyperthermia has shown promising results against breast cancer liver metastases treated with a paclitaxel-containing chemotherapy regimen^35^. Targeted hyperthermia was also shown to offer benefit in the treatment of locally advance breast cancer with alternative neoadjuvant chemotherapy regimens^36^.

It is important to finish by stating that although the proposed modelling framework has been developed based on our understanding of paclitaxel, it could potentially be applied to other molecules that freely diffuse and have high cellular retention rates. For example, drugs that bind with high affinity to abundant intracellular structures such as DNA or microtubules. This could include a wide variety of additional chemotherapeutics that are still widely used today due to their ability to achieve at least partial selectivity for tumour cells. Predicting why some patient do and don’t respond to these drugs has proved incredibly challenging, although most of those predictions have focused on genetic differences to explain the sensitivity of individual cells. If parameters relating to the 3D tumour environment are predictive instead, then these could easily have been missed due to the inherent difficulty in assessing the complex 3D tumour environment prior to treatment.

A long-term goal in cancer research has been to identify tumour specific features – the so called “hallmarks of cancer” – that can be selectively targeted with therapeutic agents^37,38^. This work has focused almost exclusively on genetic features present in cancer cells that distinguish these cells from proliferative healthy cells in the body. However, some universal features of all solid tumours that distinguish them from healthy cells in the body are their size, 3D geometry and perfusion status. If these distinguishing features of cancer were targeted by many chemotherapeutics agents in widespread use, then this could perhaps be harnessed in future to improve newer generation drugs that have less side effects on healthy cells. For example, these drugs could be modified in ways that allow them to be retained inside cells, perhaps by binding to abundant cellular structures, thus allowing them to be preferentially retained inside tumor tissue. If these drugs could then be selectively activated at a time when they are primarily retained within tumour cells, then this could increase their tumour-specificity even further, and perhaps allow otherwise toxic drugs to be used as a long-term therapy that is able to maintain tumours below a certain size threshold.

---

**Figure A1:**
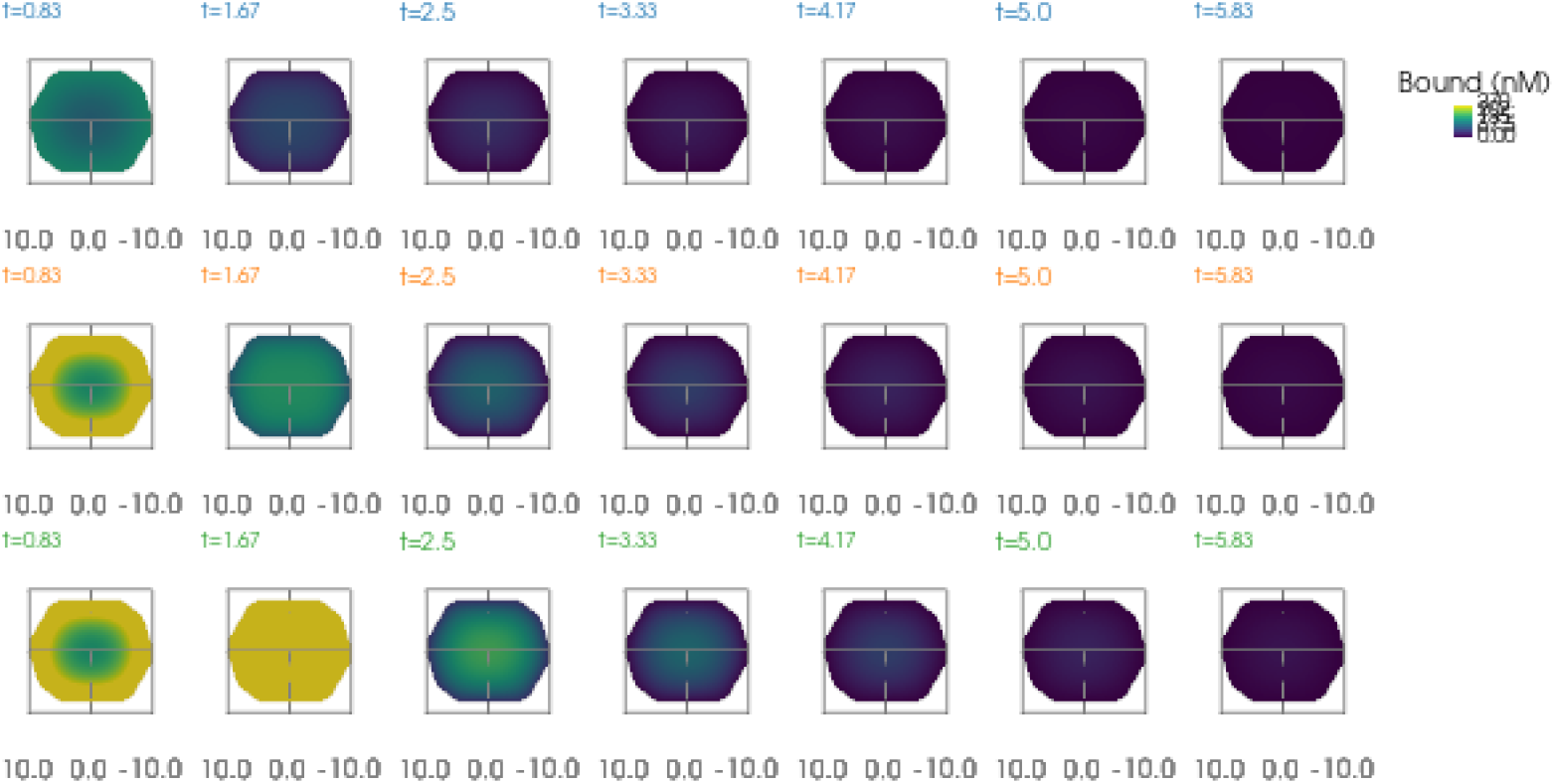
The effect of treatment duration, *t*_1_, on the penetration depth. Top row: *t*_1_ = 0.5 d. Middle row: *t*_1_ = 1.25 d. Bottom row: *t*_1_ = 2 d. Other details as in Figure 3.

## Appendix

### 3.1 Penetration dependence on treatment duration

#### 3.2 Comparing cell-based and continuum models

#### 3.3 Optimisation

#### 3.4 Slower diffusion

**Figure A2:**
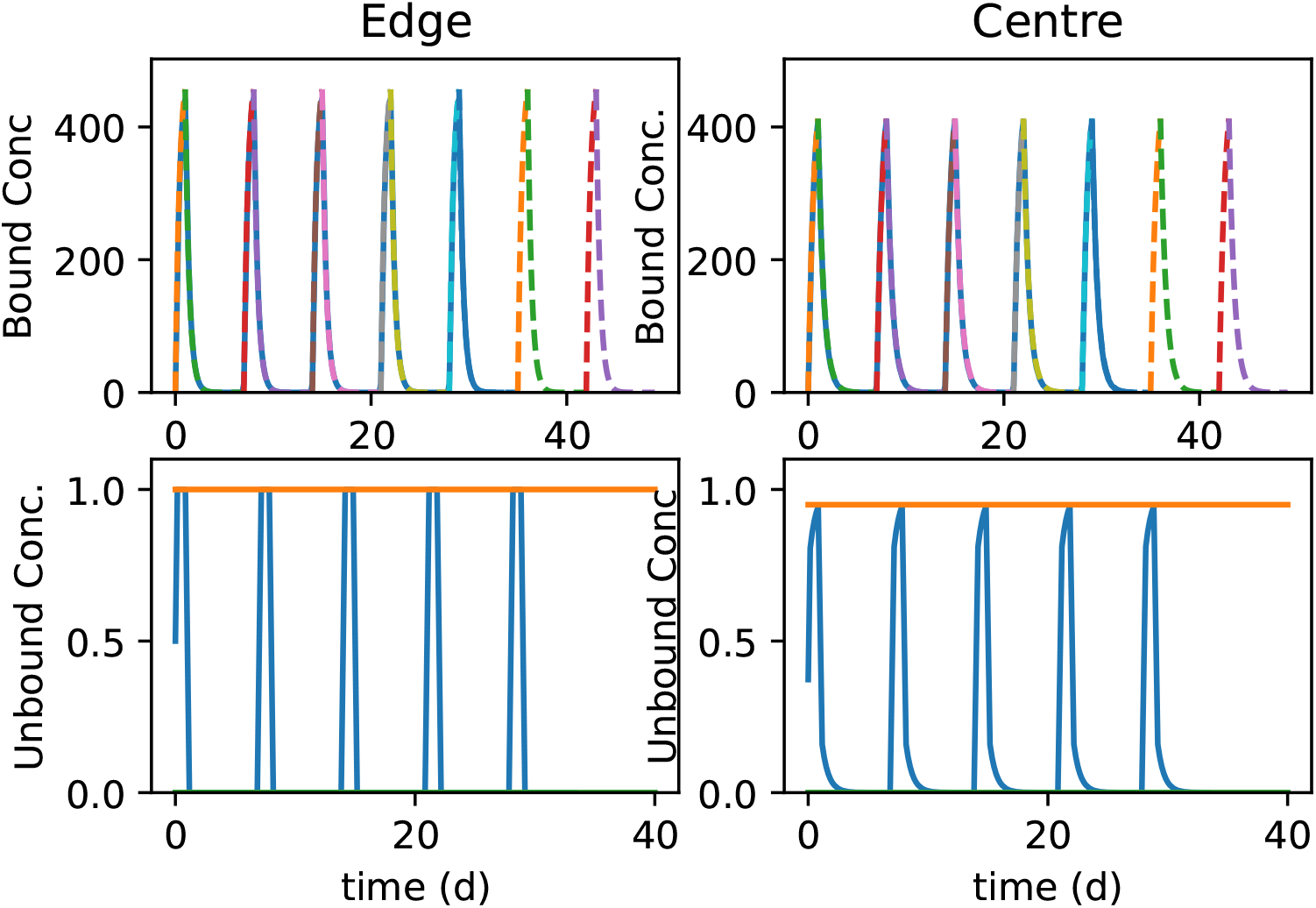
Comparing predicted paclitaxel oscillations with simulation data in small spheroids. Top row: bound intracellular paclitaxel concentrations are plotted against time in the periphery (left) and centre (right) Predicted dynamic equilibrium solution (Equation 22, dashed lines). Bottom row: unbound paclitaxel concentrations plotted against time at spheroid periphery (left) and centre (right). Solid lines: oscillation envelope. *R* = 4 c.d.

**Figure A3:**
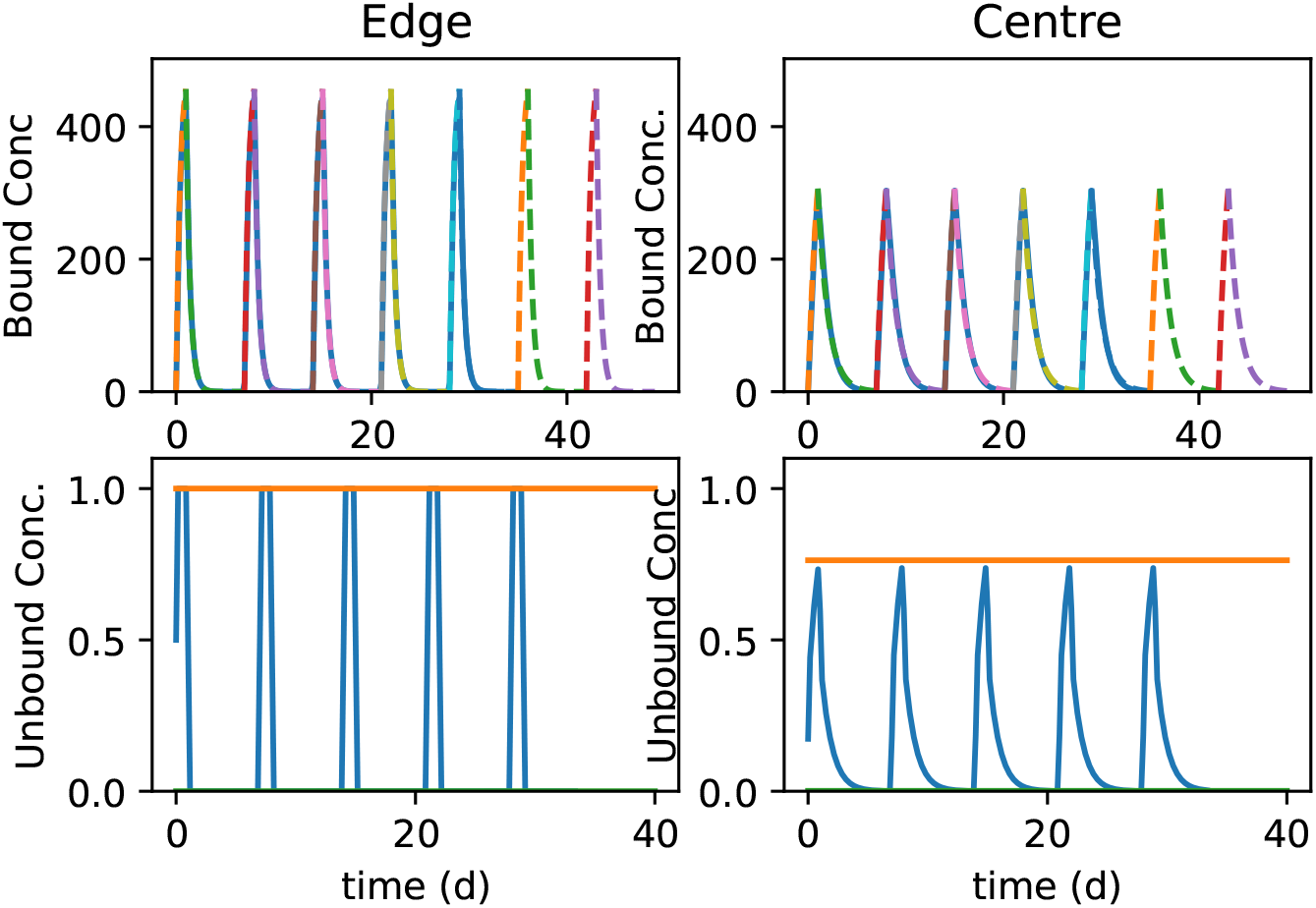
Comparing theoretical description of paclitaxel oscillations with simulation data for a larger spheroids (*R* = 8 c.d.).

**Figure A4:**
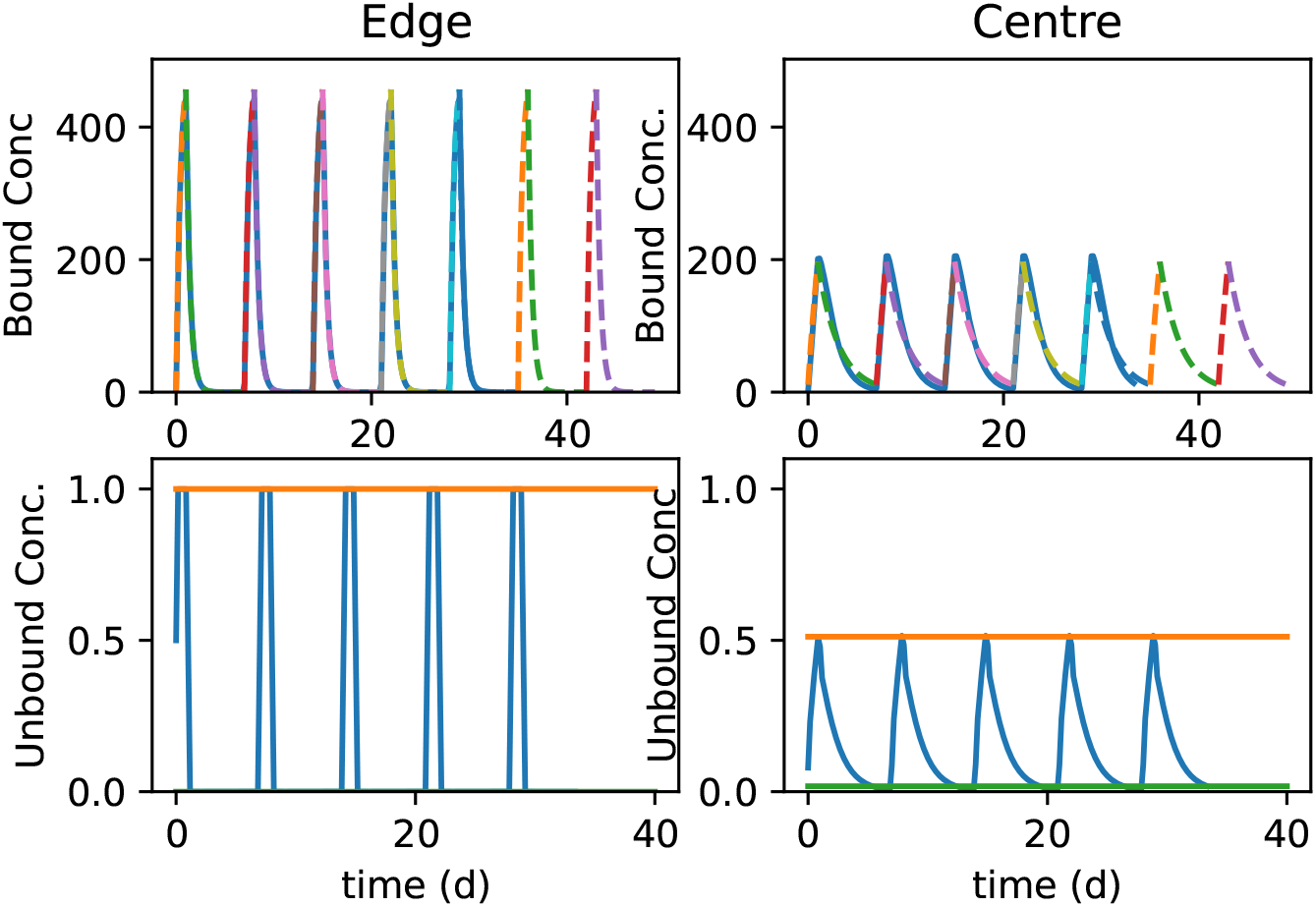
Comparing theoretical description of paclitaxel oscillations with simulation data in larger spheroids (*R* = 12 c.d.).

**Figure A5:**
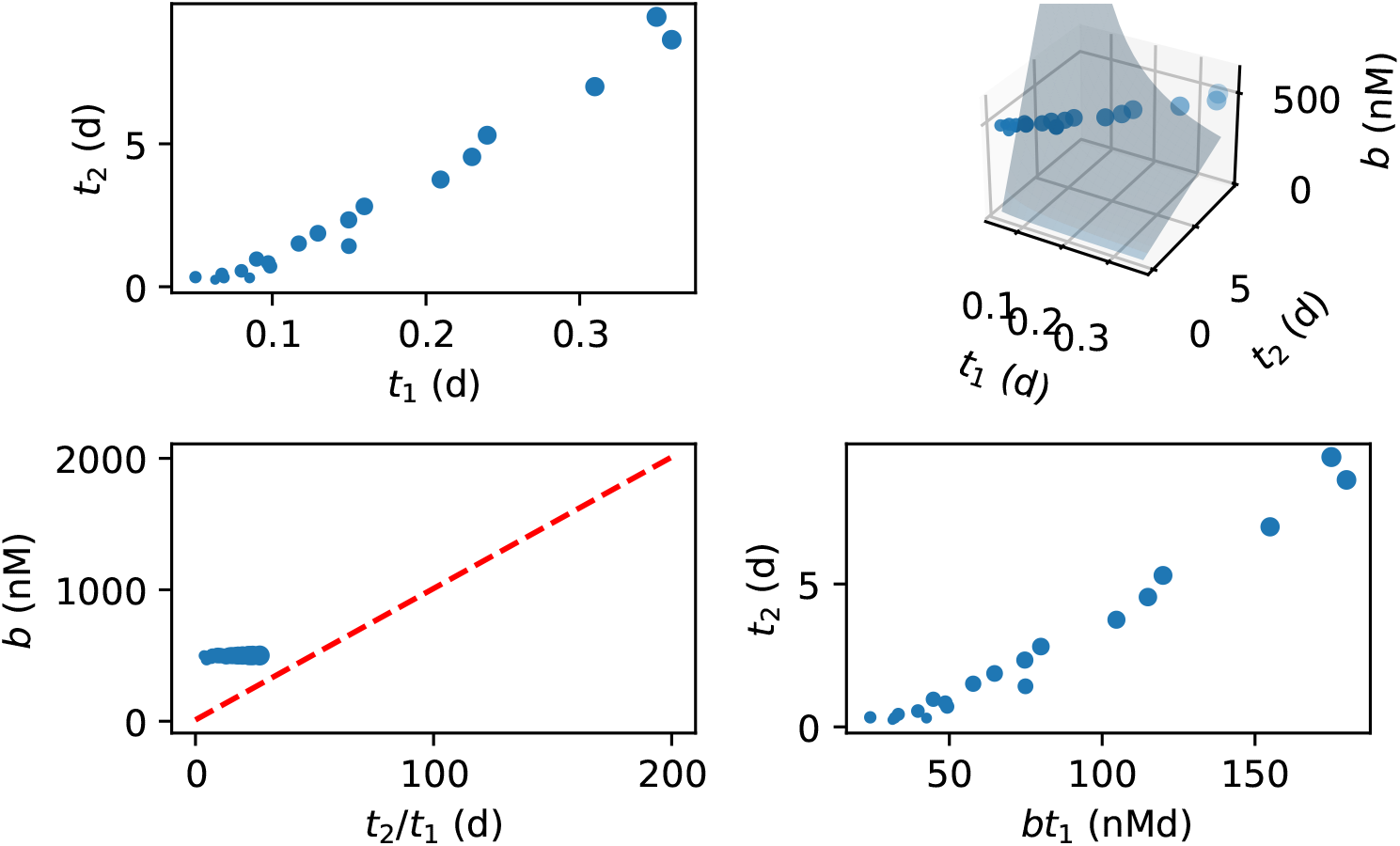
Projections of optimal parameters. (a) Projection in the *t*_1_ *− t*_2_ plane. (b) *t*_1_ *− t*_2_ *− b* space. (c) *b* is plotted against *t*_2_/*t*_1_. (d) *t*_2_ is plotted against the AUC (*bt*_1_). Other parameter values as in Table 1.

**Figure A6:**
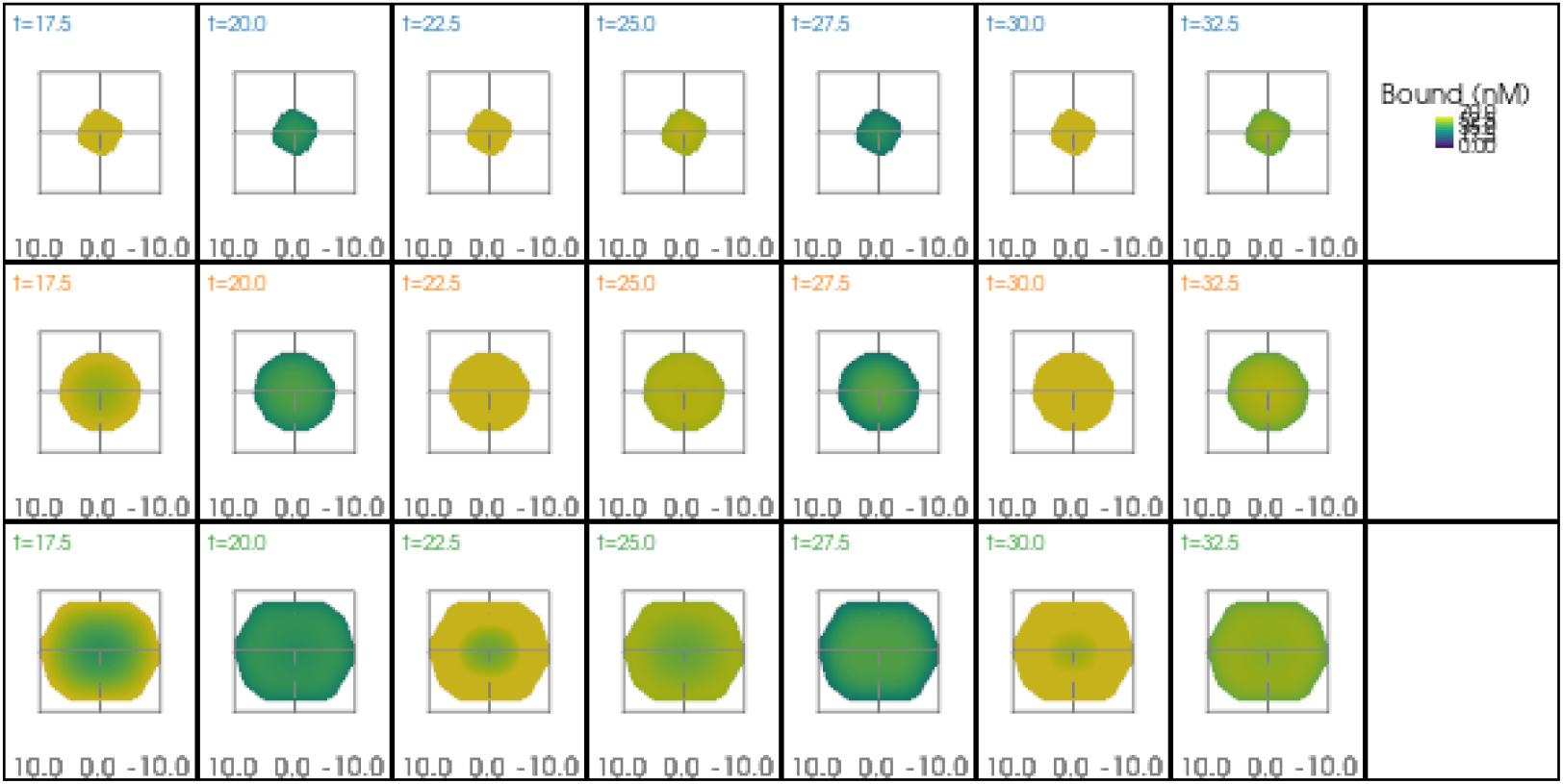
Simulation of paclitaxel uptake in spheroids of different radii with slower kinetics. Parameters *k, d* and *D* are a factor of 10 times smaller in Figure 2. Other details are the same.

**Figure A7:**
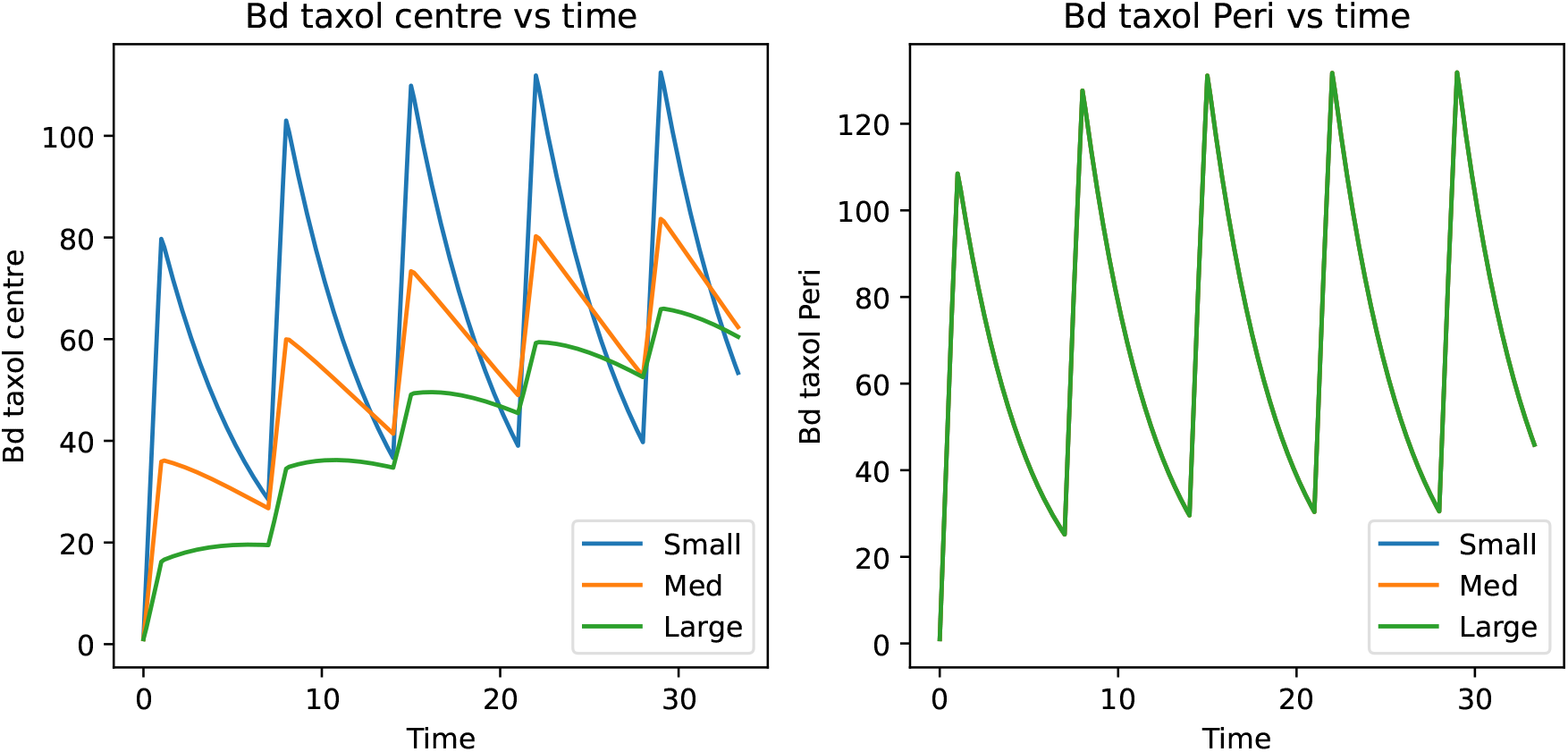
Dynamics of paclitaxel uptake in centre of spheroid depends on spheroid radius. Small, medium and large radii are 4,8 and 12 cell diameters, respectively. Other details as in Figure A6.

